# CRISPR/Cas9 targeting of MCPyV T antigen in Merkel tumors

**DOI:** 10.1101/2021.02.14.431142

**Authors:** Reety Arora, Komal Gupta, Anjali Vijaykumar

**Author notes:** **Equally contributing shared first authors** (Reety Arora, Komal Gupta). **corresponding author – Dr. Reety Arora,**. **Email ID and address**: 1. Komal Gupta.

## Abstract

Merkel cell carcinoma (MCC) is a rare, aggressive skin cancer caused either by Merkel cell polyomavirus (MCPyV) T antigen gene expression, post integration (∼80% cases), or by UV mediated DNA damage. Viral-positive Merkel tumors are not only caused by but also oncogenically addicted to tumor antigen expression. In this study we used CRISPR-Cas9 based gene-editing to develop a potential therapeutic tool for MCPyV positive MCC. CRISPR (Clustered Regularly Interspaced Short Palindromic Repeats)/Cas system is a genome editing technology whereby a guide RNA (gRNA) molecule, targets a Cas endonuclease to a specific genomic site, using sequence homology, and induces a double strand break. To target MCPyV T antigens, we designed a strategy using 2 gRNAs targeting the T antigen genomic region that would cut off a substantial portion of the gene thereby rendering it dysfunctional. We validated the MCPYV T antigen targeting efficiency of our gRNAs, both individually and together by *in vitro* cleavage assays. Finally, to translate this finding, we delivered this CRISPR system in patient-derived MCC cell lines and show reduction in T antigen gene expression. Our proof-of-concept study shows that 2 MCPyV targeting CRISPR/Cas gRNAs in combination can knock out MCPyV T antigen, thus, being of therapeutic importance. We hope that this CRISPR system can be potentially delivered *in vivo* for advancing MCPyV positive MCC treatment in the future.

## INTRODUCTION

Merkel Cell Carcinoma (MCC) is a rare and aggressive neuroendocrine cancer of the skin^1^. Prolonged UV exposure, advanced age and loss of immune competence are risk factors for this cancer^2,3^. It is either caused by Merkel cell polyomavirus (MCPyV)^4^ or long term Ultraviolet (UV) exposure^5-7^. In the majority of human population MCPyV is a harmless, asymptomatic and life-long infection. However, MCPyV initiates an aggressive cancer if it gets integrated into the susceptible host genome and acquires viral mutations that lead to replication incompetence^4^. In tumors, MCPyV expresses 2 oncoproteins: the small and large Tumor antigens^3,4^. These T antigens drive tumorigenesis by targeting various tumor suppressors and regulating transcription^8-10^.

A majority of MCC cases have been found to be MCPyV positive (∼60-80%)^1,3,4^. In virally caused cancers the T antigens are the major drivers in tumor development. Interestingly, T antigen knockdown in MCC cell lines initiates cell cycle arrest and cell death^9,11^. As MCPyV positive MCC tumors are oncogenically addicted to the expression of T antigens, they are attractive targets for therapeutics.

In this study we developed a CRISPR Cas9^12^ system to knock-down MCPyV T antigen expression. This is the first report to use CRISPR Cas9 to reduce MCPyV T antigen expression and we hope that with further validation and analysis this will serve as a potential targeted gene therapy system in the future.

The CRISPR (Clustered Regularly Interspaced Short Palindromic Repeats)/Cas9 system is a programmable gene editing system adapted from the bacterial immune system^12^. CRISPR uses a guide RNA (gRNA) molecule that targets a Cas endonuclease to a specific genomic site using sequence homology and PAM (Protospacer adjacent motif) recognition^13-15^. Thus, the end result of Cas9-mediated DNA targeting is a double stranded break (DSB) within the target. This break is then corrected by the cell, usually via the mutagenic Non homologous end joining (NHEJ) pathway, resulting in disruption of target gene^16^. Using transfection or lentiviral transduction, it is possible to induce the cleavage of specific sequences in the human genome by the expression of SpCas9 and a sgRNA molecule. Gene inactivation using this system is highly efficient, and SpCas9 has been previously used to target various genes including HIV proteins and E6/E7 oncoproteins in cervical carcinoma^17-19^.

Here we report the use of the CRISPR SpCas9 system to target the T antigen oncogenes in MCPyV positive MCC cell lines. First, we designed several gRNAs and tested their ability to target T antigen expression. Then, we designed a strategy to use 2 validated gRNAs and measured their efficiency using an *in vitro* cleavage assays. Further, we used these two gRNAs together to target the T antigen loci in cells expressing MCPyV T antigens and Cas9. Our study is the first to report the use of the CRISPR-Cas9 system for targeting MCPyV T antigen genes in Merkel tumors.

## MATERIALS AND METHODS

### sgRNA design

13 gRNAs were designed to target the small and large Tumor antigen genes of MCPyV. Three CRISPR design tools: Broad institute software (broadinstitute.org/gpp/public/analysis-tools/sgrna-design), CRISPR-MIT (http://crispr.mit.edu) and CRISPOR were used. MKL-1 (Genbank Accession #: FJ173815.1) and MS-1 (Genbank Accession #: JX045709.1) MCPyV sequences were used as target DNA sequences. gRNAs with highest specificity score and with lowest possible off targets were selected. Synthetic gRNA oligonucleotides were cloned into pLentiCRISPRv2 (Addgene #52961) at BsmBI (# R0580S, NEB) restriction sites. Plasmid DNA was prepared using Stbl3 competent cells (#C7373-03, Invitrogen) and Qiagen midi prep kits (#12143).

### Synthesis of sgRNAs using IVT

Two oligonucleotides were designed as DNA templates, per gRNA, for *in vitro* transcription (IVT). Forward oligo for Cas9 gRNA synthesis consisted of the T7 promoter (TAATACGACTCACTATAGG), guide RNA, and start of Cas9 gRNA scaffold. Reverse oligo consisted of complete gRNA scaffold with complementarity to 15nt in the forward oligo (shown below). Three G’s were added at the gRNA start for efficient T7 transcription.

Target substrate for C3 gRNA *in vitro* cleavage (<u>TARGET</u> / pam):

5’ GATGAAAGCTGCTTTCAAAAGAAGCTGCTT<u>AAAGCATCACCCTGATAAAG</u>gggGAAATCCTGTTAT 3’

crRNA oligo for C3 gRNA *in vitro* transcription (**T7** / ADDED G / <u>GUIDE</u> / *SCAFFOLD):*

Forward primer: **TAATACGACTCACTATA**GGG<u>AAAGCATCACCCTGATAAAG</u>*GTTTTAGAGCTAGAA* Reverse primer: *AAAAGCACCGACTCGGTGCCACTTTTTCAAGTTGATAACGGACTAGCCTTATTTTAA CTTGCTATTTCTAGCTCTAAAAC*

Target substrate for C13 *in vitro* cleavage (<u>TARGET</u> / pam):

5’ CCAGTGTACCTAGAAATTC<u>TTCCAGAACGGATGGCACCT</u>gggAGGATCTCTTCTGCGATGAATC 3’

crRNA oligo for C13 gRNA *in vitro* transcription (**T7** / ADDED G / <u>GUIDE</u> / *SCAFFOLD):*

Forward primer: **TAATACGACTCACTATA**GGG<u>TTCCAGAACGGATGGCACCT</u>*GTTTTAGAGCTAGAA* Reverse primer: *AAAAGCACCGACTCGGTGCCACTTTTTCAAGTTGATAACGGACTAGCCTTATTTTAA CTTGCTATTTCTAGCTCTAAAAC*

Templates for IVT were amplified by the following PCR reaction: Denaturation at 95°C for 3 mins followed by 30 cycles of 95°C – 30s, 45°C – 30s and 72°C for 30s and final annealing at 72°C for 7 min. The PCR products were cleaned up using PCR clean up kit (# A9282, Promega). sgRNAs were synthesized using the MEGAShortScript™ Kit (# AM1354, Invitrogen) following the manufacturer’s protocol with 2ug PCR product template over night at 37°C. The RNA was treated with 1ul of DNase TURBO for 15 minutes followed by purification using the MEGAclear Transcription Clean-Up Kit (# AM1908, Invitrogen).

### *In vitro* cleavage assay

*In vitro* cleavage reaction was performed with purified SpCas9 protein (500ng) (Kind gift from Dr. Praveen Vemula’s Laboratory) at 37°C in cleavage buffer consisting of 20 mM HEPES (pH 7.5), 150 mM KCl, 10 mM MgCl2, 1% glycerol and 0.5mM DTT for 1 hour. 300 ng of sgRNA and 200 ng of the target DNA were used for the reaction. PCR amplicons of sT and LT gene regions from plasmid RAZ1 (Addgene #114381) were used as target DNA templates (Refer Supplementary Table 1 for primers). The PCR amplicon was purified using Ampure beads (#A63881, Beckman Coulter) as per manufacturer’s protocol. The IVC reactions were stopped by treatment with 2ul of Proteinase K (10mg/ml) at 55°C for 30 minute. 1x gel loading dye was added to the reactions and samples were run on 2% agarose gels (# RM273, HIMEDIA).

### Plasmid Construction

We cloned the NCCR and T antigen derived from MKL-1 into a pLentiCMV backbone vector We used the target region PCR amplicon as the DNA substrate.

### Gibson Cloning the gRNAs in pDECKO mCherry

For expressing the gRNAs in MCPyV positive cell lines, C3 and C13 were cloned into pDECKO mCherry lentiviral plasmid using Gibson assembly. The backbone plasmid (kind gift of Dr. Debjyoti Chakroborty, IGIB) was digested using BsmBI at 55°C for 2 hours. The cut plasmid was run on a 0.8% agarose gel and the linear band was gel extracted. The oligonucleotides for insert 1 consisting of the gRNAs, T7 and part of H1 promoter and gRNA scaffold were designed using http://crispeta.crg.eu. Insert 1 was cloned into digested vector and was transformed into Stbl 3 competent cells. Positive clones were selected using colony PCR followed by plasmid isolation. This intermediate plasmid was again digested by BsmB1. The constant insert 2 consisting of remaining part of gRNA 1 scaffold and H1 promoter was PCR amplified from the complete pDECKO plasmid and was cloned into the intermediate plasmid. The reaction was transformed in Stbl 3 competent cells. Positive clones were validated using colony PCR and sequencing.

### Cell culture conditions

HEK 293T cells were obtained from ATCC and grown in Dulbecco’s Modified Eagle Medium (DMEM) supplemented with 10% Fetal Bovine Serum (FBS) 1X Penicillin/Streptomycin (Pen Strep, catalog # 15140122). Merkel Cell Carcinoma cell lines, MKL-1 were obtained from ECACC (#09111801) and cultured in Roswell Park Memorial Institute (RPMI) Media with 10% FBS and 1X Penicillin/Streptomycin. The cells were grown at 37°C with 5% CO_2_.

### Lentiviral vector construction and transduction

Lentiviral vectors were constructed by co-transfecting HEK 293 T cells with plasmid of interest, the Gag-Pol packaging plasmid psPAX2 (Addgene #12259), and pVSVG (Addgene #8454) expressing plasmid in a ratio of 5:3.75:1.25. Lipofectamine® LTX (Invitrogen #15338100) was used in 3:1 ratio with the total amount of DNA to be transfected. DNA, Lipofectamine LTX (Invitrogen #15338500) and OptiMEM mix were incubated at RT for 20 minutes and added to cells in antibiotics free media. Viral media was collected 48h and 72h post transfection. The viral media was filtered through 0.45 μm filter followed by concentration through Amicon column and addition of 1/3^rd^ volume of LentiX Concentrator (Takara #631231). Viral pellet was obtained by spin down at 1500g at 4°C, resuspended in RPMI media and added to the MKL-1 cells to be transduced. Screening of gRNAs was done by transducing pLentiCRISPRv2 plasmid containing the gRNAs in MKL-1 cells followed by puromycin (1μg/mL) selection. pLentiCRISPRv2 (Addgene #52961) without gRNA was used for mock transduction. For constructing MKL-1 pDECKO mCherry C3+C13 cell line, MKL-1 cells were transduced with pDECKO mCherry C3+C13 plasmid. pDECKO mCherry empty vector (EV; without gRNAs) was used as an empty vector for mock transduction. MKL-1 Cas9 cell line was developed using pCWCas9 Blast (Addgene #83481) with Blasticidin (10ug/mL) selection for 10 days post transduction. Cas9 was expressed by 1ug/ml of Doxycycline induction.

### Western blot analysis

Cells were lysed in EBC buffer (50 mM Tris pH-8, 150 mM NaCl, 0.5mM EDTA, β mercaptoethanol-1:10,000, 0.5% NP-40, 1 mM PMSF supplemented with protease inhibitor cocktail (Roche, Cat #118575). Protein lysate was quantitated using Pierce BCA protein assay kit. The lysate was denatured in 1x Laemmli buffer by boiling at 99°C for 5 minutes. The proteins were separated using SDS-PAGE and transferred to a polyvinylidene fluoride (PVDF) membrane (#88518, Biorad) The PVDF membrane was blocked with 5% Blotto (SC-2325, Santacruz) in 1X TBS-T. The blot was developed with primary antibodies overnight at 4°C a s follows: anti-MCPyV Tumor antigen monoclonal antibody (1:5000;Ab3 and Ab5 were used (put citation)), anti-Cas9 monoclonal antibody (1:1000; Cell signaling #14697), and anti-Vinculin monoclonal antibody (1:10000; Sigma #V9131). The PVDF membrane was incubated in peroxidase-labeled secondary antibodies. The ECL™ Prime Western Blotting System (#RPN2232, GE Healthcare) was used for the detection of chemiluminescence according to the manufacturer’s protocol. The chemiluminescence was detected using ImageQuant™ Las 4000 (GE Healthcare).

## RESULTS

To target the integrated MCPyV Tumour antigens present in MCPyV positive MCC cells, we designed 13 sgRNAs. These gRNAs are complementary to regions between 247 to 1254 nucleotides of MKL-1 genomic sequence that encompass both LT and sT (Table 1 and Supplementary Fig 1). 5 gRNAs (C1 – C5) target Exon 1 (common region between sT and LT), 4 gRNAs (C6 – C9) target only sT and the rest of the 4 gRNAs (C10 – C13) target only LT. These sgRNAs were cloned into pLentiCRISPRv2 which also expressed SpCas9 protein under the control of a U6 promoter. To investigate the cleavage activity of these 13 sgRNAs, we transduced MKL-1 cells with these 13 constructs. Immunoblotting showed significant reduction in the expression of both ST and LT by 2 gRNAs – C3 and C13 (Supplementary Figure 1). We thereby proceeded with these two gRNAs for further experimentation (Figure 1).

**Table 1.**
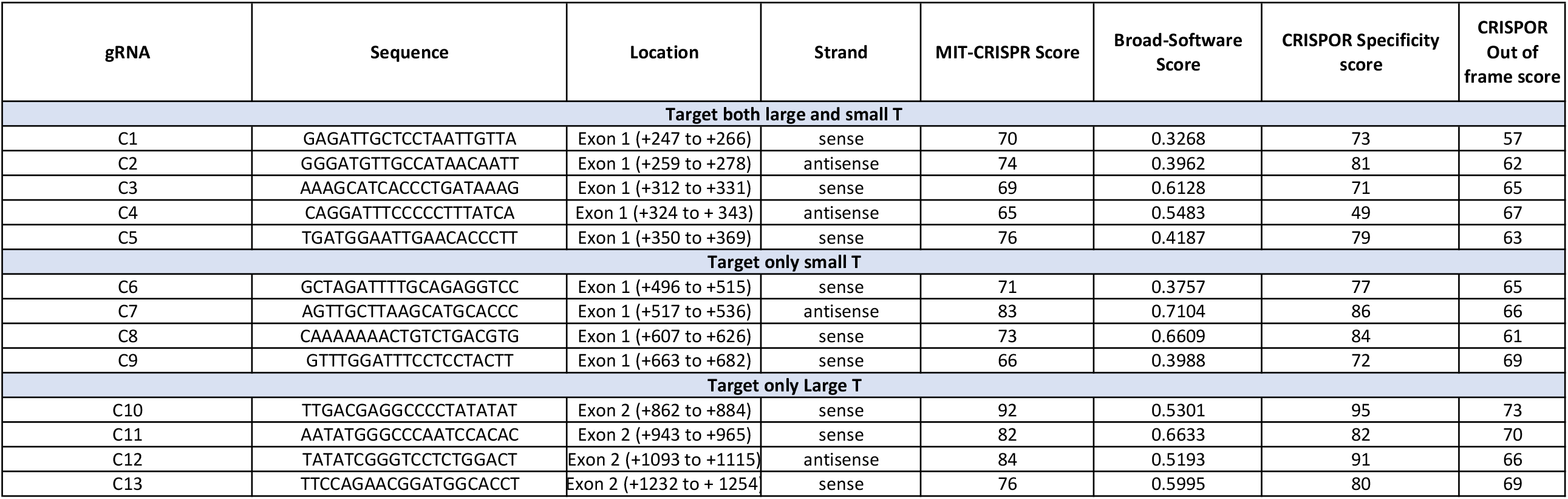
gRNAs targeting MCV T antigen genomic region.

**Figure 1.**
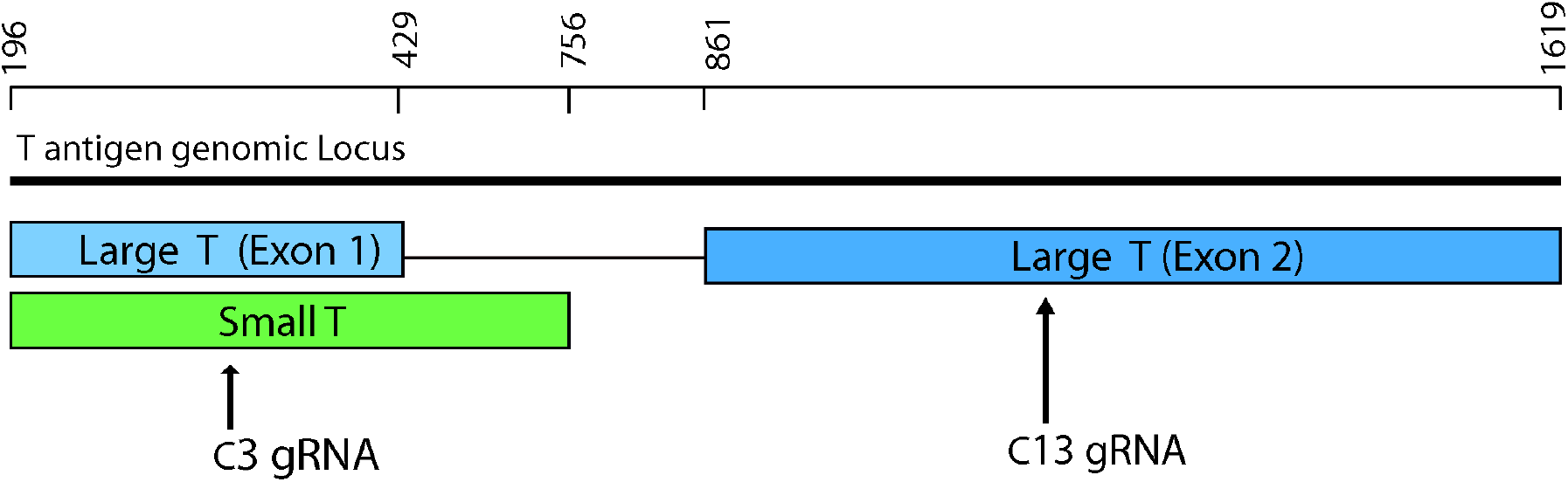
gRNA annotation on MCPyV T Antigens. gRNA C3 targets the genomic region encoding both the exon 1 of Large T antigen and small T antigen and gRNA C13 targets the genomic region targeting exon 2 of large T antigen. A cut at both sites causes an excision of the T antigen genomic region shown.

To further validate the cleavage efficiency of gRNAs C3 and C13, we performed *in vitro* cleavage assays (Figure 2). We cloned the NCCR and T antigen derived from MKL-1 into a pLentiCMV backbone vector and used the target region PCR amplicon as the DNA substrate for our assays. The two gRNAs efficiently cleaved the target substrate as shown by the cut bands of sizes 562bp, 367bp and 195bp for C3 and 512bp and 400bp for C13, respectively (Figure 2). As shown, we used two different SpCas9 enzymes as technical replicates to test our gRNAs. To address the time taken for this cleavage, we further performed a time course assay. Both the gRNAs were able to cleave the target within 10 minutes (Figure 3). Therefore, C3 and C13 can effectively target and cleave MCPyV T antigen genomic sequence.

**Figure 2.**
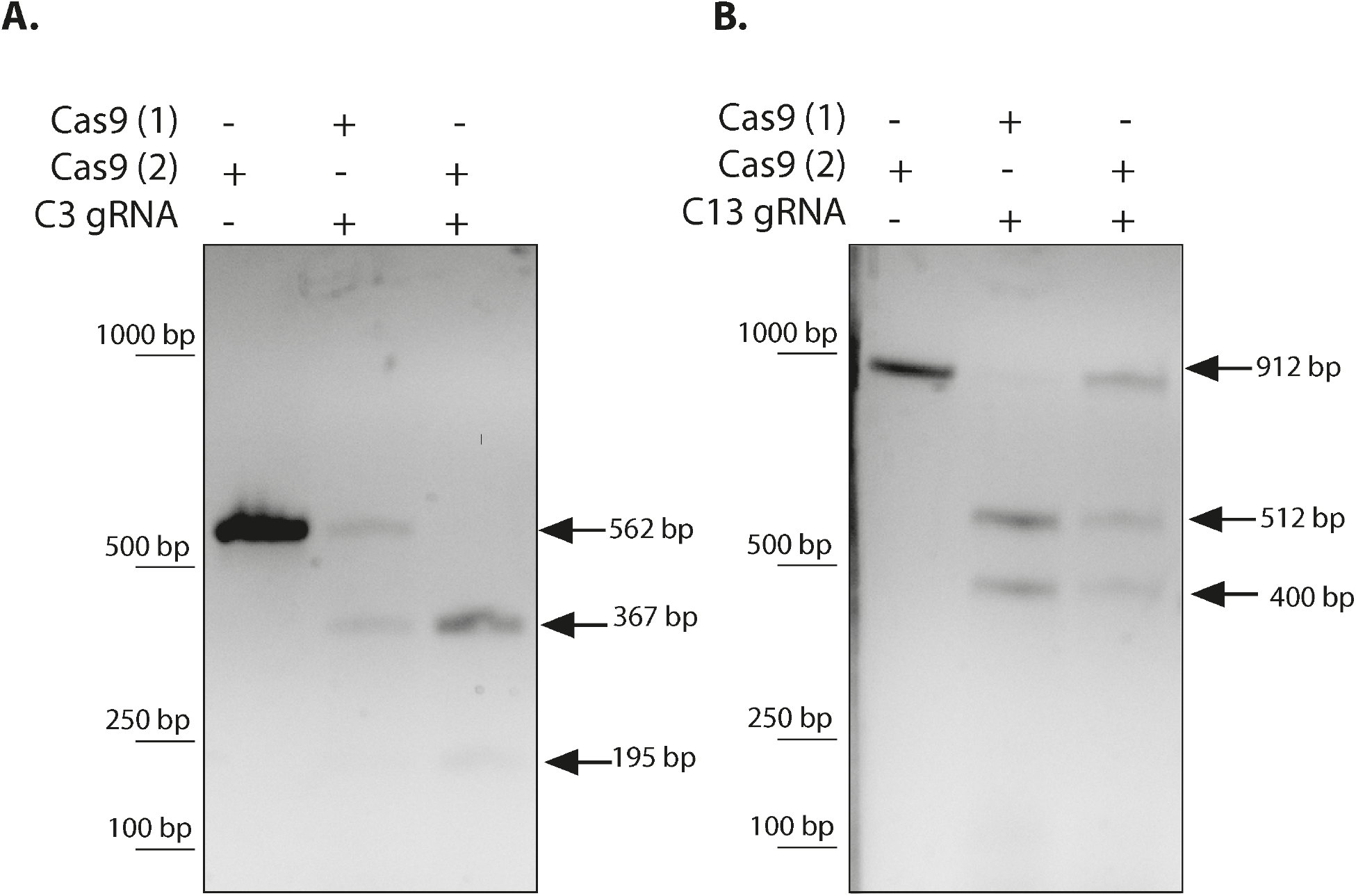
*In vitro* Cleavage Assay for gRNAs C3 and C13. The MCPyV C3 and C13 gRNA target region template was amplified from RAZ2 plasmid via PCR. *In vitro* cleavage reaction performed on this template shows SpCas9 mediated MCV target cleavage in the presence of gRNAs **(A**.) C3 and **(B**.) C13 as indicated by arrows.

**Figure 3.**
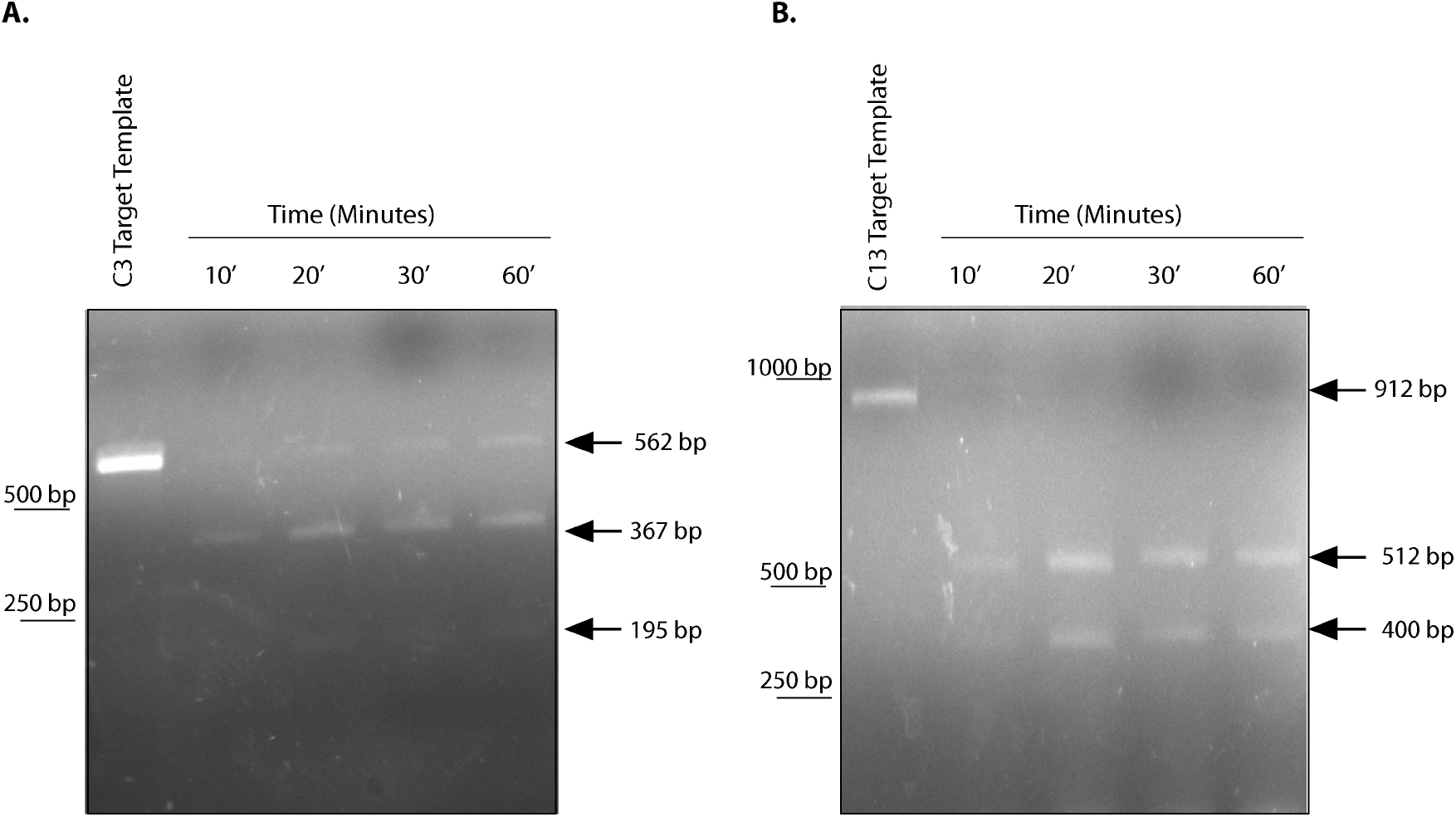
Time course *In vitro* cleavage assays for gRNAs C3 and C13. SpCas9 can cleave the MCV target template within 10 minutes.

Next, using *in vitro* cleavage assays, we validated the combinatorial target cleavage by gRNAs C3 and C13 (Figure 4). The effective cleavage activity is shown by the 3 cleaved bands at 1131 bp, 940 bp and 374 bp as indicated by the arrows (Figure 4).

**Figure 4.**
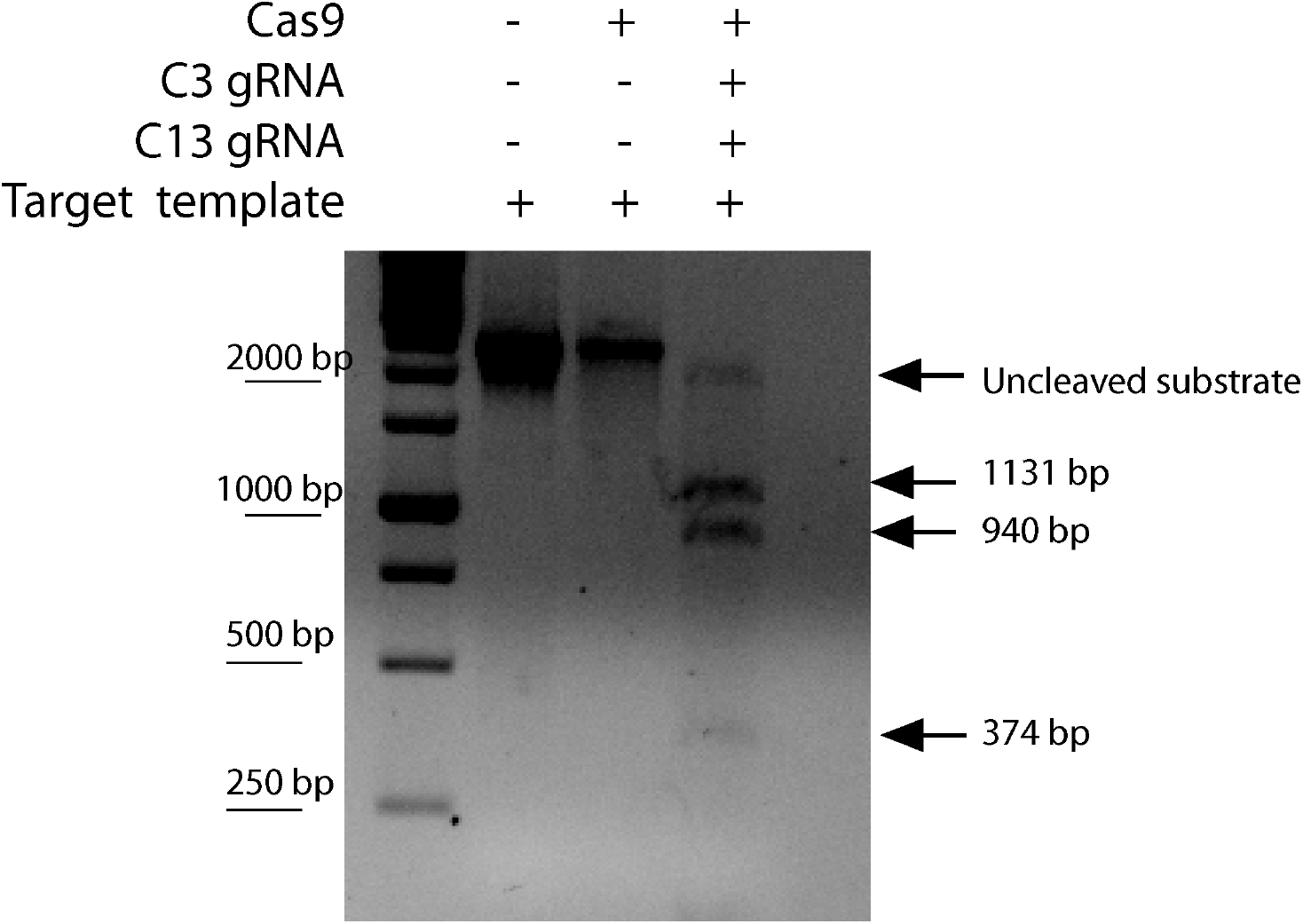
*In vitro* Cleavage Assay for gRNAs C3 and C13 together. A 2445 bp long MCV target was amplified from RAZ2 plasmid. Cas9 in the presence of C3 and C13 gRNAs led to a successful combinatorial target cleavage.

To investigate the effects of these CRISPR/Cas9 constructs in combination, we cloned C3 and C13 into pDECKO mCherry vector. This vector expresses 2 gRNAs together under U6 and H1 promoters respectively (Figure 5A).

**Figure 5.**
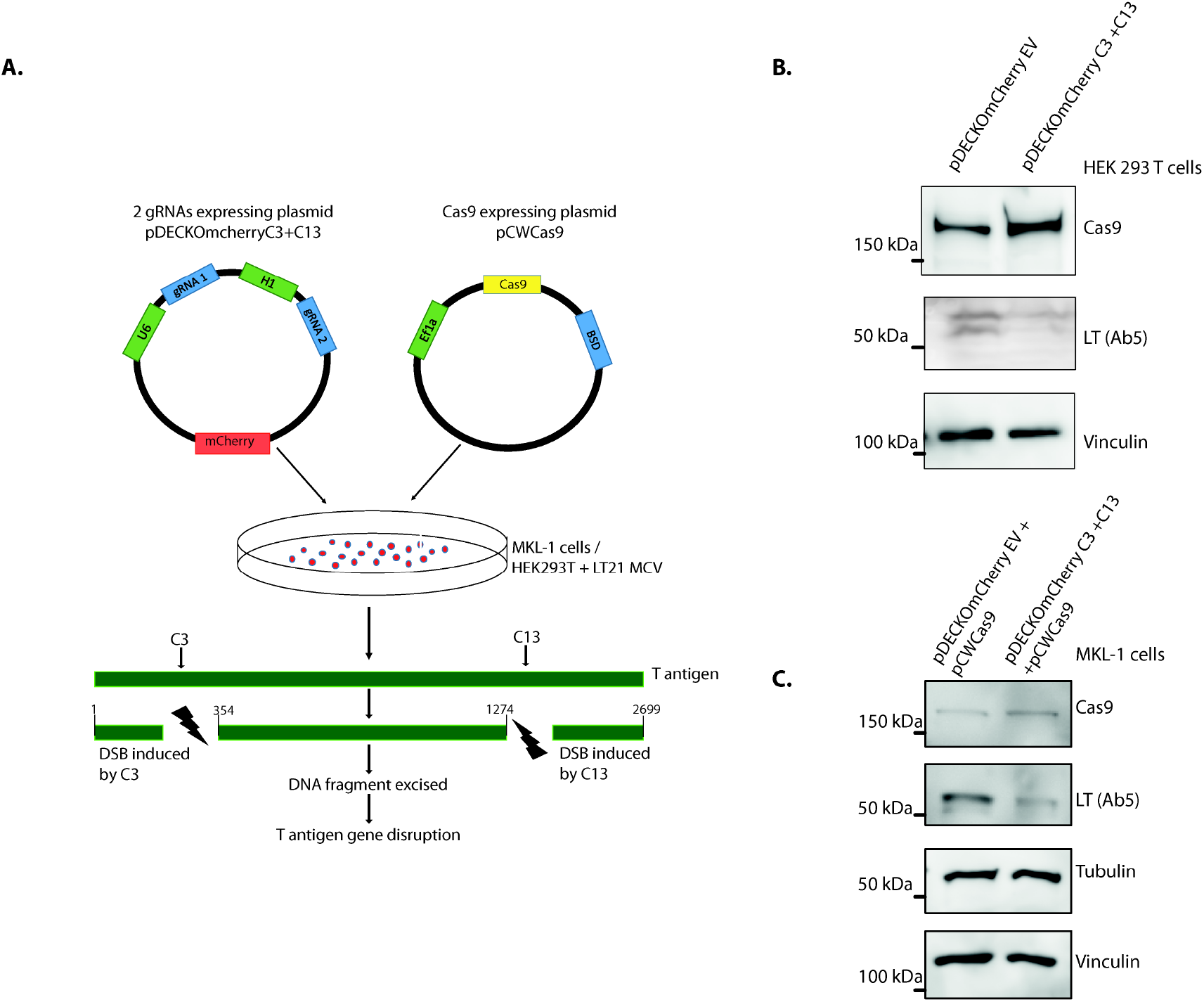
CRISPR Cas9 Targeting of viral protein in MCPyV T antigen expressing cells. **(A**.) Schematic showing strategy for gRNA and Cas9 expression. **(B**.) Immunoblot showing T antigen targeting in HEK293T cells expressing both MCPyV T antigen and Cas9. **(C**.) Immunoblots showing T antigen loss in MKL1 cells.

To test the efficiency of this construct we co-transfected HEK293T cells with Merkel cell polyomavirus (MCC LT162) early region expressing plasmid, Cas9-expressing plasmid and the pDECKO mCherry empty vector (EV) or pDECKO mCherry C3 + C13. Western blot data revealed a loss of LT expression upon transfection of pDECKO C3 + C13 relative to pDECKO mCherry empty vector (Figure 5B).

Further, we targeted MCPyV T antigen locus in MKL-1 cells using C3 and C13 gRNAs using lentiviral-vector delivery. We transduced MKL-1 cells and harvested cells post-selection. We first developed stable pDECKO mCherry EV and pDECKO mCherry C3+C13 expressing cell lines. For this we infected MKL-1 cells with pDECKO lentiviral vectors followed by Puro selection. Additionally, we expressed Cas9 using a Tet-inducible system in these MKL-1 stable cell lines using Blasticidin selection. Infection followed by selection of MKL-1 cells with Cas9 and 2 gRNAs showed a significant reduction in MCPyV T antigen expression (Figure 5C).

## DISCUSSION

Changes in DNA causes cancer, and in virally caused cancers such as MCC this change refers to the integration of a viral genome. Scientists have been searching for easy ways to correct DNA changes linked to cancer and other diseases for ages, when CRISPR came along in 2013.

The gene-editing tool CRISPR is a game changer offering a precise, customizable, fast and an easy-to-use pair of molecular scissors to cut human DNA within cells.

For MCPyV positive Merkel cell carcinoma, RNAi has already been used to interrogate the effect of blocking MCPyV T antigen gene expression in cancer cells and testing phenotype^9^. Knocking down of T antigens via shRNA kill MCC cancer cell lines. The oncogenic dependency of Merkel tumor cells on T antigen expression renders them essential for their survival, thus making them excellent targets for potential therapy.

Our strategy used CRISPR to target the MCPyV T antigen genomic region and cause two double stranded DNA cuts. The excision of an important DNA sequence of the viral genome region, flanking between the two target sites, by DSBs at two sites caused a disruption in gene expression. This strategy is more robust as compared to targeting via single gRNA as the excision of DNA sequence leads to knock out of target gene. Further, we validated the gRNAs in HEK293 cells stably expressing MCPyV T antigens and also in MCPyV positive MKL-1 cells and found a significant reduction in MCPyV T antigen levels.

CRISPR using Cas9 has several advantages to RNAi; besides being more precise, it targets the genomic DNA as opposed to mRNA and causes a permanent gene disruption and thus a stronger effect.

The field of CRISPR is growing rapidly since its discovery and its application in human gene editing was established. Since we began the study, Cas13a and Cas13d have been used successfully and efficiently to target protein-coding mRNA transcripts to knock-down gene expression in human cells^20-22^. This Type VI CRISPR Cas system holds great promise for the future of gene expression loss mediated therapy^23^.

Our study provides a proof of principle for the use of CRISPR technology to edit out MCPyV T antigen from the Merkel tumor genome. Removal of MCPyV T antigens by using the SpCas9/sgRNA combinations system has significant clinical potential in the treatment of MCPyV positive MCC. However slipping CRISPR components into lab-grown cells in one thing, the delivery of the Cas enzymes and gRNA into a cancer patient’s body remains the challenge.

In the future, this tool can be delivered *in vivo* using Adeno associated virus (AAV) vectors^24^, which are smaller and safer than lentivirus or retrovirus vectors and can infect with high titers. As AAV vectors have also been employed previously for gene therapy they could be potential vectors for *in vivo* delivery of Cas9 and gRNAs^24^.

Recently Chen et al., 2019 developed a promising alternative CRISPR delivery system using nanocapsules^25^. Their 25 nm diameter polymer capsule can carry Cas9 and a gRNA. Using specific peptides and other ingredients, this capsule can target predetermined cell types. It is highly stable in the bloodstream and releases its contents once inside the cell^25^.

Researchers are continuing to explore different ways to fine-tune this delivery of CRISPR components to specific organs and cell-types of the human body^24,26,27^. We hope that in the near future, such a smooth delivery will enable the two gRNAs described, to be used for targeting MCPyV for MCC therapy successfully prolonging patient lives.

## ACKNOWLEDGEMENTS

We are grateful to Dr. Jingwei Cheng and Dr. James Decpario for valuable discussions and guidance. We are also very grateful to Dr. Debojyoti Chakraborty and Mohammed Azhar from IGIB, Delhi for guidance and advice on CRISPR related experiments. Special thank you to Lamiya Dohadwala and Bhavana Nayer for technical assistance and to Prof. Sanjeev Galande for scientific comments. RA would also like to thank Zaina, Ziyah, and Karan for their love, strength, and support.

---

**Figure S1.**
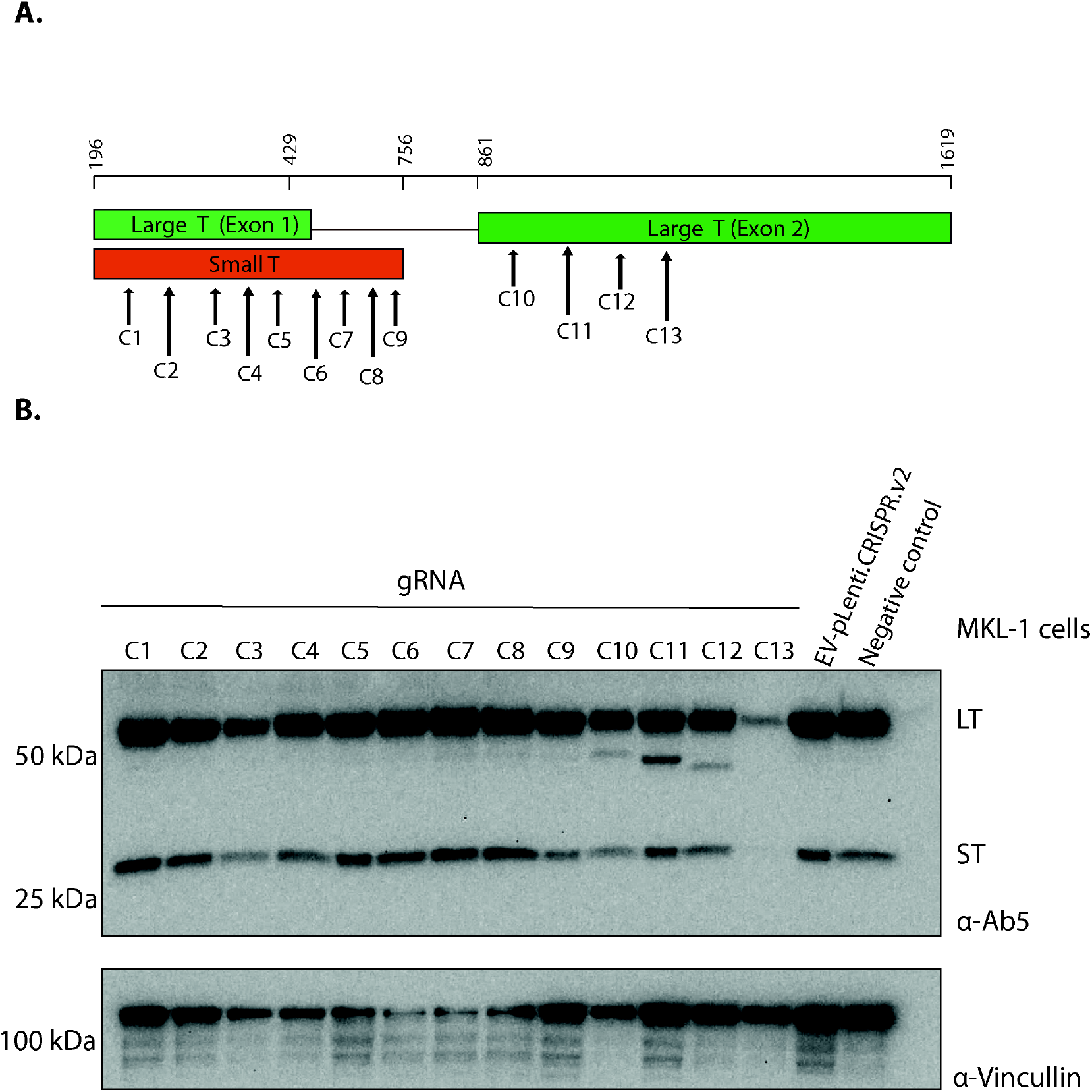
gRNAs target MCV T antigen. **(A**.) 13 gRNAs were designed to target MCPyV T antigen. **(B**.**)To** investigate the cleavage activity of these 13 sgRNAs, MKL-1 cells was transduced with 13 gRNA plasmid constructs. lmmunoblotting above shows significant reduction in the expression of both ST and LT by 2 gRNAs - C3 and Cl 3. Abs was used to measure MCPyVT antigen expression and Vincullin was used as a housekeeping control. The empty vector plenti.CRISPR.v2 and only MKL-1 cell lysate was used as negative controls.

**Supplementary Table 1.**
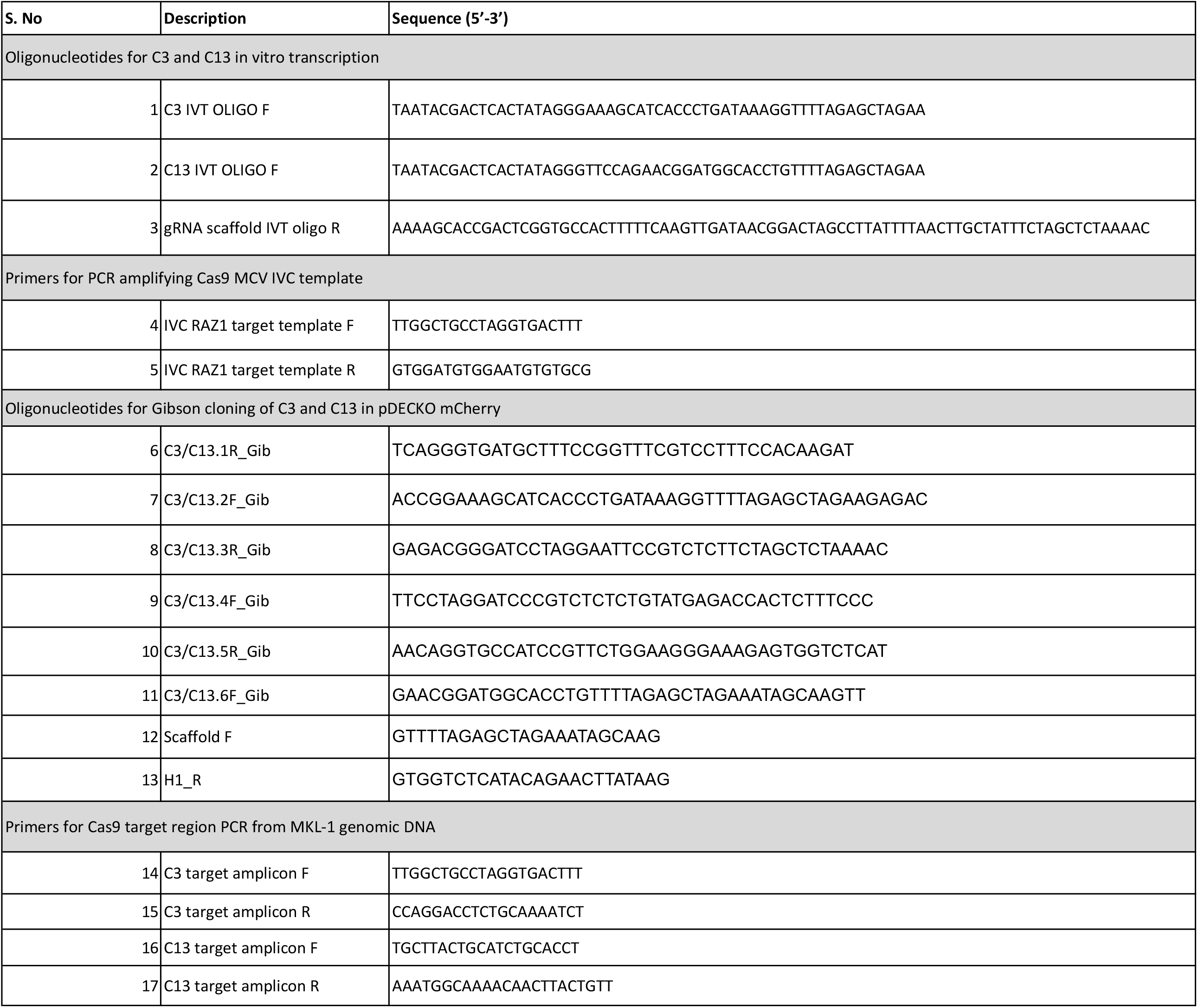
Primers Used.

